# Chloroplast Genome Data of *Tecticornia indica*, a Hygrohalophytic C4 Plant Species

**DOI:** 10.1101/2023.12.29.573661

**Authors:** Richard M. Sharpe, Bruce Williamson-Benavides, Gerald E. Edwards, Amit Dhingra

## Abstract

*Tecticornia indica* is a hygrohalophytic, NAD-malic enzyme-type C4 plant species with near-aphyllous photosynthetic shoots. Total cellular DNA was extracted from the aphyllous shoots and branches of *Tecticornia indica* for subsequent High Throughput Sequencing (HTS). Illumina HiSeq 2000 technology produced a total of 91,825,666 reads. Reads were trimmed and filtered to retain base calls corresponding to a 99.9% base call accuracy. A *de novo* assembly produced 413,729 contigs and 45,238 scaffolds with a maximum length of 120,548 nucleotides, an N_50_ of 881, and an average depth of coverage of 16.29 nucleotides. A subset of 2,196,299 reads were used for assembling the chloroplast genome at an average depth of coverage of 1,094 nucleotides. Sanger sequencing was performed to validate the LSC-IR-SSC junctions and to complete the missing regions of the chloroplast genome. The assembled chloroplast genome map and *T. indica* genomic sequence is expected to facilitate genetics and genomic research focused on understanding function of genes involved in halophytic and drought tolerant traits.

**SPECIFICATIONS TABLE:** 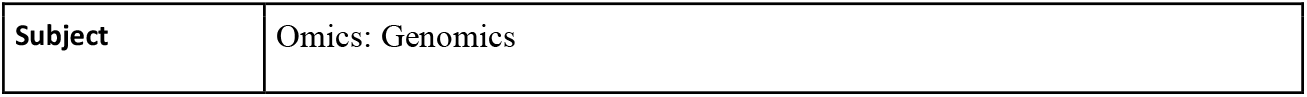

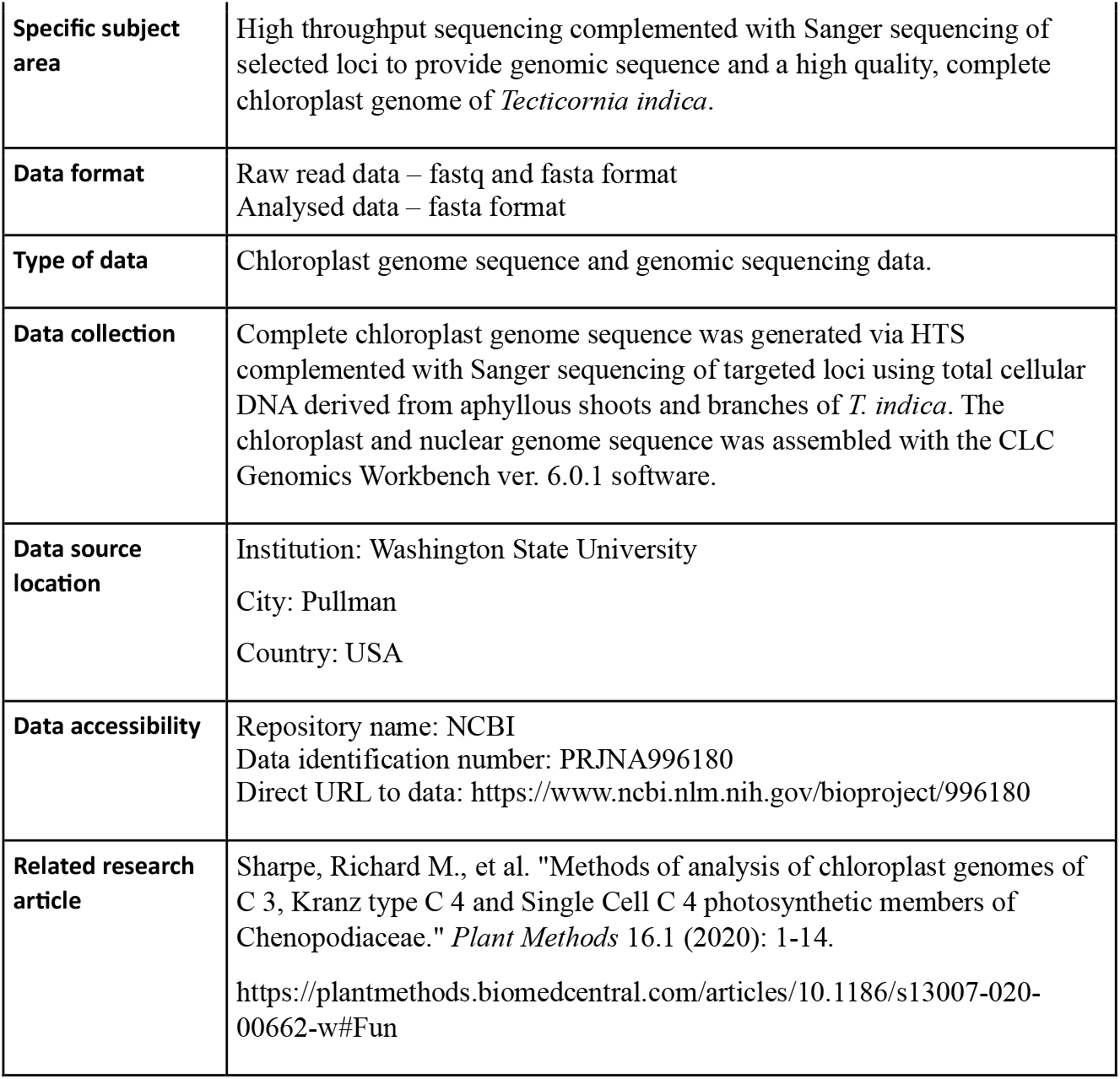

**VALUE OF THE DATA:**

- Genomic sequence information of *Tecticornia indica* and its chloroplast map will be useful for genomics research.
- Genomics and genetics scientists, bioinformatics and biotechnologists in the fields of photosynthesis, taxonomy and drought tolerance will benefit from these data.
- These data can be used in developing molecular markers and quantitative trait loci in identification and selection of halophyte and drought tolerance, taxonomic resolution, and variables in photosynthetic genes.

## BACKGROUND

Our research group has worked on the sequencing and releasing of chloroplast genomes of dicot species that include representative species having C3 and C4 -type photosynthesis. The motivation was to generate the chloroplast sequence of *Tecticornia indica*, (C4 Plant Species), a genome that had not been published yet. The release of representative C3 and C4 dicot species can help determine whether the chloroplast genomes between C3 and C4 species are highly conserved (in size and composition), and the degree of difference between the species.

## DATA DESCRIPTION

### DNA sequencing, validation, and contig assembly

Illumina sequencing of *T. indica* total cellular DNA resulted in a total of 91,825,666 reads. Of these, a total of 2,196,299 reads, with a mean read length of 76.03, were used for the assembly of the *T. indica* chloroplast genome. The average, minimum, and maximum coverage was 1,094, 6, and 76,450, respectively.

Three contigs of 89,774nt, 18,747nt, and 10,540nt length and high homology to the large single copy (LSC), small single copy (SSC), and inverted repeat (IRA & IRB) regions of the *B. sinuspersici* chloroplast genome, were obtained (GenBank accession no. KU726550). In comparison to the *B. sinuspersici* chloroplast genome, the *T. indica* genome was missing undefined length of regions at the IRB-LSC and LSC-IRA junctions, a 6,581nt section in the LSC contiguous to the IRB, a region of 14,269nt, from which a section of 11,951nt was originally assembled to the LSC, and a section of 2318nt in the IR. When combined, these generated an IR with a total length of 24,809nt.

Missing regions in *T. indica* were successfully amplified through primer walking PCR and subsequent amplicon sequencing (Supplementary Information Table S1). The repeat content (>8nt) did not show any major differences between the missing and contiguous regions. Most of the assembled chloroplast genome presented a coverage above 40x. There were a total of 11 sections with coverage below 40x (Supplementary Information Table S2). These genomic areas were validated with the use of Sanger sequencing. After DNA sequencing, assembly, and validation, the assembled chloroplast genome map showed the canonical quadripartite organization as well as highly conserved gene sequences associated with the majority of higher land plants.

### Size, organization, gene and repeat content of the chloroplast genome

The complete chloroplast genome of *T. indica* assembled to a size of 152,701nt. The chloroplast genome included a pair of IR regions of 24,809nt in length separated by a SSC region of 18,701nt, and a LSC region of 84,382nt (Figure 1). GC content of the chloroplast genome was similar to what was previously reported in other species within Chenopodiaceae [1]. The entire plastome, LSC, SSC and IRA and IRB showed a 34.1%, 29.4%, 42.9%, and 42.8% GC content respectively as well as a 36.4% overall GC content. *T. indica* chloroplast genome contained a total of 122 genes, from which 98 were unique and 22 were duplicated. There were 29 distinct tRNAs identified, from which 12 were duplicated in the IRs. The chloroplast genome sequence consisted of 61% coding region represented by 54% of protein coding genes and 7% of RNA genes.

**Figure 1.**
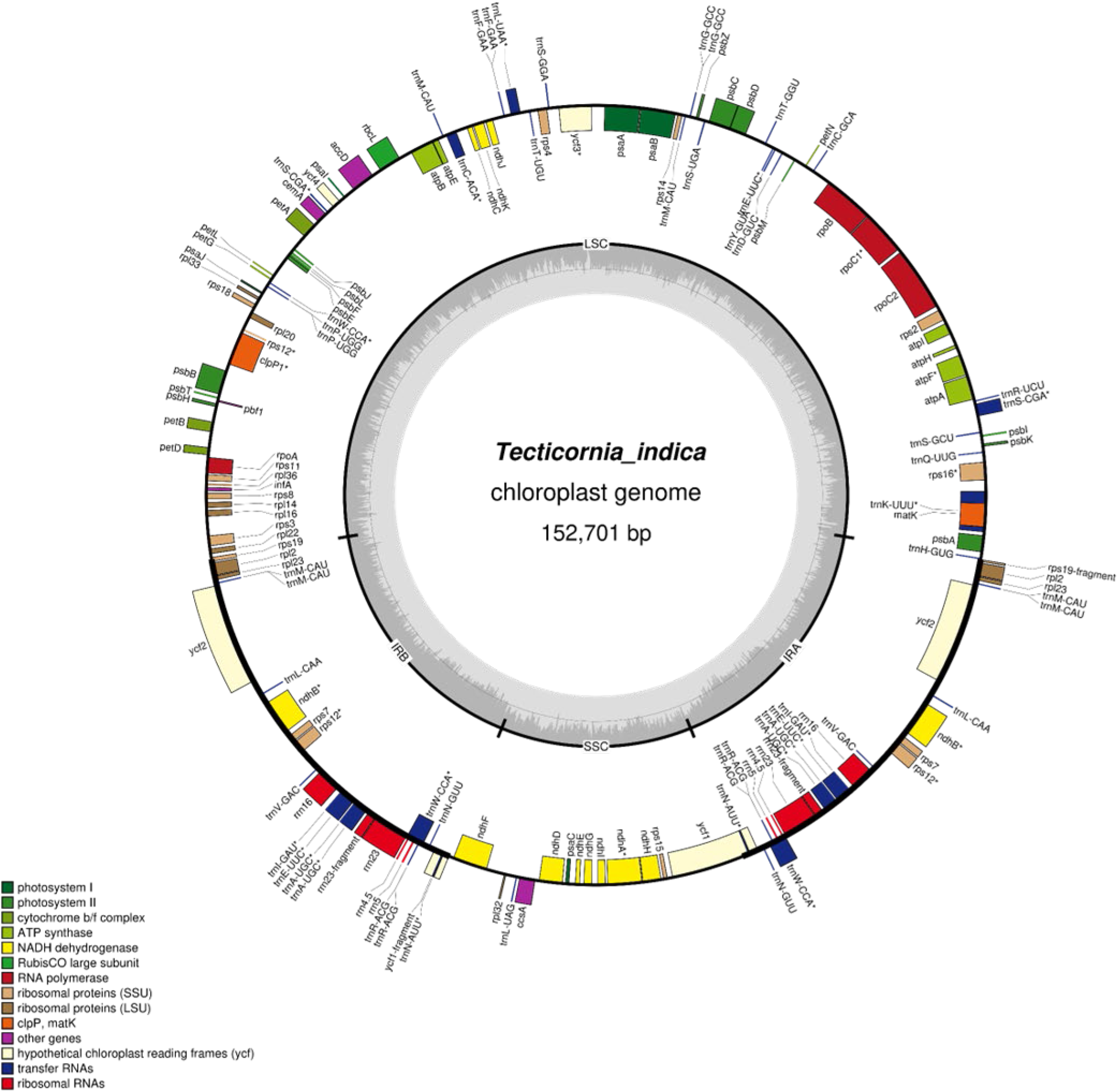
Physical Map of *Tecticornia indica* chloroplast genome. Sequence information is available in the Supplementary Information File 4.

Gene content and order of the *T. indica* chloroplast genome is similar to the previously reported chloroplast genomes within Chenopodiaceae [1]. A small portion (1.018 kb) of the YCF1 gene (5.622 kb) was duplicated in the IR. A similar pattern has been previously reported for *V. vinifera, S. oleracea* and *B. vulgaris* as well as other Chenopodiaceae species such as A *retroflexus, B. muricata, B. cycloptera, B. sinuspersici, S. aralocaspica*, and *S. maritima* [1].

The *T. indica* chloroplast genome had 40 repeats ranging from 30 to 44nt (Table 1). The majority of these repeats, 34 repeats, were between 30 and 40 bp in length similar to other Chenopodiaceae species [refer to Table 4 in Sharpe et al., (2020)]. The number of repeats in genes and intergenic regions were similar to other Chenopodiaceae species [1]. The presence of repeats varied for *ycf*1, *ycf*2, and *psa*A genes, as for most of the species in the Chenopodiaceae family. However, *T. indica* did not have any repeats in the ycf3 introns. Interestingly, *T. indica* had six repeats in the *psa*A gene, which is the highest number of repeats within the Chenopodiaceae family. A total 0f 67 microsatellites were identified, of which the majority, 51, represent mononucleotide repeat microsatellites. A total of 6, 3, and 7, tetra-, penta- and compound-mononucleotide repeat microsatellites, respectively, were identified (Supplementary Information Table S3).

**Table 1.** Total number of repeats distributed by length and location in *Tecticornia indica* chloroplast genome.

|  |  | Total number of repeats | (%) |
| --- | --- | --- | --- |
| Length (nt) | 30-34 | 25 | 61.0 |
|  | 35-39 | 9 | 22.0 |
|  | 40-44 | 5 | 12.2 |
|  | 45-54 | 1 | 2.4 |
|  | 55+ | 1 | 2.4 |
| Location | Intergenic | 16 | 39.0 |
|  | Exons | 25 | 61.0 |
|  | Introns | 0 | 0.0 |

## EXPERIMENTAL DESIGN, MATERIALS AND METHODS

### Plant Material and DNA extraction

*Tecticornia indica* plants were grown in greenhouse conditions with a 14/10 h photoperiod, light regime of 120 PPFD, and average day/night temperatures of 28°C/18°C. Total cellular DNA was isolated from tissue derived from aphyllous shoots and branches of *T. indica* tissue with a Urea Lysis Buffer Method.

### DNA sequencing, validation, and contig assembly

DNA sequencing, validation and contig assembly was performed as detailed by Sharpe et al., (2020). Briefly, a paired-end library was generated and sequenced on the Illumina HiSeq 2000 utilizing the 100PE chemistry. Quality control on raw sequence data, read trimming, and filtering out for low quality reads was performed using CLC Genomics Workbench ver. 6.0.1 (CLC) (QIAGEN, Redwood City, CA, USA). Mapping of reads was conducted in CLC using the following mapping parameters: mismatch cost 2, insertion cost 3, deletion cost 3, length fraction 0.8 and similarity fraction 0.9.

BLASTN searches on NCBI (https://www.Ncbi.nlm.nih.gov/) were performed to identify contigs with high homology to chloroplast large single copy (LSC), small single copy (SSC) and inverted repeat (IR). The identified IR contig was reverse complimented and the overlapping borders of each of the identified contigs were aligned to assemble the *T. indica* chloroplast genome sequence in the following order of LSC+IR+SSC+IR.

Chloroplast contig junctions were PCR amplified using flanking primers (Supplementary Information) and were sequenced using Sanger sequencing (Eurofins Genomics, KY). Overlapping PCR fragments ranged in size from 0.2 to 0.5 kb. Primer walking and Sanger sequencing were utilized to identify missing regions in the LSC+IRA and IRB+LSC junctions (Supplementary Information). Raw sequence data are accessible through the NCBI database under the BioProject number PRJNA996180 (https://www.ncbi.nlm.nih.gov/bioproject/?term=PRJNA996180).

### Size, organization, and gene and repeat content of chloroplast genome

*T. indica* chloroplast genome was annotated and visualized with GeSeq [2] which incorporates the Dual Organellar Genome Annotator (DOGMA) [3], and OrganellarGenomeDRAW (OGDRAW) [4].

Pair-wise comparisons were performed to analyze gene order and content for the *T. indica* chloroplast genome. These comparisons included eight Chenopodiaceae chloroplast genomes [1] as well as three chloroplast reference genomes *V. vinifera* (NC_007957.1), *S. oleracea* (AJ400848.1), and *B. vulgaris* (EF534108.1).

Repeat identity and structure was obtained using REPuter [5]. A minimum repeat size of 30 bp and a Hamming distance of 3 (>90% sequence identity) was the set parameter. Microsatellites were identified with MISA software [6] using a minimum stretch of 10 for mono-, six for di-, five for tri-, and three for tetra-, penta-, and hexa-nucleotide repeats, and a minimum distance of 100 nucleotides between compound microsatellites.

## LIMITATIONS

Not applicable

## ETHICS

### STATEMENT

All authors have read and follow the ethical requirements for publication in Data in Brief and confirming that the current work does not involve human subjects, animal experiments, or any data collected from social media platforms.

### CRediT AUTHOR STATEMENT

Richard M. Sharpe: Investigation, formal analysis, Data curation, Writing - Original Draft Bruce Williamson-Benavides: Investigation, formal analysis, Data curation, Writing - Original Draft, Visualization. Gerald E. Edwards: Conceptualization, Resources, Funding acquisition, Writing - Review & Editing. Amit Dhingra: Conceptualization, Methodology, Resources, Writing - Review & Editing, Supervision, Funding acquisition

## ACKNOWLEDGEMENTS

This work was funded in part by a National Science Foundation Grant MCB 1146928 to GEE and AD, and Washington State University Agriculture Center Research Hatch Grant WNP00011 and startup funds from Texas A&M AgriLife Research and Texas A&M University to AD. BWB acknowledges graduate research assistantship support from Washington State University Graduate School.

